# Alopecia areata susceptibility variant identified by MHC risk haplotype sequencing reproduces symptomatic patched hair loss in mice

**DOI:** 10.1101/308197

**Authors:** Akira Oka, Atsushi Takagi, Etsuko Komiyama, Shuhei Mano, Kazuyoshi Hosomichi, Shingo Suzuki, Nami Motosugi, Tomomi Hatanaka, Minoru Kimura, Mahoko Takahashi Ueda, So Nakagawa, Hiromi Miura, Masato Ohtsuka, Yuko Haida, Masayuki Tanaka, Tomoyoshi Komiyama, Asako Otomo, Shinji Hadano, Tomotaka Mabuchi, Stephan Beck, Hidetoshi Inoko, Shigaku Ikeda

**Author notes:** To whom correspondence should be addressed. or, Akira Oka, Atsushi Takagi, Etsuko Komiyama, Shuhei Mano, Kazuyoshi Hosomichi, Shingo Suzuki, Nami Motosugi, Tomomi Hatanaka, Minoru Kimura, Mahoko Takahashi Ueda, So Nakagawa, Hiromi Miura, Masato Ohtsuka, Yuko Haida, Masayuki Tanaka, Tomoyoshi Komiyama, Asako Otomo, Shinji Hadano, Tomotaka Mabuchi, Stephan Beck, Hidetoshi Inoko, Shigaku Ikeda.

## Abstract

**Background:** Alopecia areata (AA) is a highly heritable multifactorial and complex disease. However, no convincing susceptibility gene has yet been pinpointed in the major histocompatibility complex (MHC), a region in the human genome known to be associated with AA as compared to other regions.

**Results:** By sequencing MHC risk haplotypes, we identified a variant (rs142986308, p.Arg587Trp) in the coiled-coil alpha-helical rod protein 1 (*CCHCR1*) gene as the only non-synonymous variant in the AA risk haplotype. Using CRISPR/Cas9 for allele-specific genome editing, we then phenocopied AA symptomatic patched hair loss in mice engineered to carry the *Cchcr1* risk allele. Skin biopsies of these alopecic mice showed strong up-regulation of hair-related genes, including hair keratin and keratin-associated proteins (KRTAPs). Using transcriptomics findings, we further identified CCHCR1 as a novel component of hair shafts and cuticles in areas where the engineered alopecic mice displayed fragile and impaired hair.

**Conclusions:** These results suggest an alternative mechanism for the aetiology of AA based on aberrant keratinization, in addition to generally well-known autoimmune events.

## Background

Alopecia areata (AA) is a complex disease defined by focal or universal hair loss, most commonly occurring on the scalp[1]. Although benign, AA can seriously impair quality of life, such as negative effects on self-perception and self-esteem[2], and even attempted suicide in rare cases[3]. The prevalence of AA is approximately 0.1% to 0.2% in the United States, with a lifetime risk of 1.7%[4], while a twin study suggested a 55% concordance rate in identical twins with a significant occurrence of AA in families[5]. In addition, the prevalence rate of AA in families has been shown to be higher than that in the general public, though the rate varied in each study and population examined[6–10].

AA is driven by cytotoxic T lymphocytes and was found to be reversed by Janus kinase (JAK) inhibition in model mice by clinical treatment[11], demonstrating that the immunological pathway is a factor in this multifactorial disease. Previous genome-wide association study results have implicated a number of immune and non-immune loci in the aetiology of AA[12–14], though none has yet been demonstrated to be causative for the disease. Thus, no variants among those have provided experimental evidence for biological functions between alleles and AA pathogenesis. Alleles of the human leukocyte antigen (HLA) genes within the major histocompatibility complex (MHC) region on chromosome 6p21.3 have thus far shown the strongest associations with AA among the human genome[12–14] in observations of different ethnic groups. However, the strongest associations with AA have not been supported by functional evidence.

The genetic architecture of the MHC region shows that multiple haplotypes with the highest degree of diversity are often maintained in a population by balancing selection, and that positive selection can occasionally generate long-range haplotypes[15, 16]. The strong linkage disequilibrium (LD) observed in such haplotypes can mask the ability to discriminate between a *bona fide* variant associated with disease and a variant influenced by LD. This limitation can be addressed by analysis of microsatellites that have higher mutation rates than SNPs, thus leading to breakup of apparently invariant SNP haplotypes into lower frequency haplotypes for functional analysis[17]. Analysis of multi-allelic microsatellites may therefore be an effective strategy for identifying rare disease-associated haplotypes in the MHC.

With this background in mind, we implemented a 4-step study design. First, we performed association analysis using microsatellites for the entire MHC region with AA patients and healthy controls to identify risk haplotypes associated with AA. Second, we sequenced representative risk and control haplotypes to identify variants that were present only in identical risk haplotypes based on all of the variants detected. Third, we performed whole exome sequencing of the variant-associated candidate gene, haplotype estimation, and extended haplotype homozygosity (EHH) analysis in all subjects. Finally, we engineered mice carrying the human risk allele using allele-specific genome editing with the CRISPR/Cas9 system and performed functional evaluations.

## Results

### Association analysis and risk haplotype sequencing

A total of 171 AA patients and 560 healthy controls were enrolled for association analysis using 22 microsatellites spanning the human leukocyte antigen (HLA) class I and II regions (chr6: 30407655-32854116, hg19), including the *HLA-C* locus, which was previously implicated to be involved in AA[18]. We detected a single microsatellite, *D6S2811* (allele *208*, OR = 3.41, CI 95% = 1.94-5.99, *P* = 3.39 × 10^−5^), which was shown to be significantly associated with AA after Bonferroni correction (Fig. 1a **and Additional file1: Table S1 and S2**). Pair-wise evaluation of these multi-allelic loci indicated that a strong long-range LD was maintained across the assayed region of the MHC (**Additional file1: Figure S1**). Furthermore, estimation of haplotypes in 3 loci from *D6S2811* to *D6S2930* showed that the risk haplotype was defined as the segment that displayed a significant association with AA (MShap01, OR = 3.78, CI 95% = 2.00 – 7.16, *P* = 6.57 × 10^−5^) (Table 1).

**Fig. 1.**
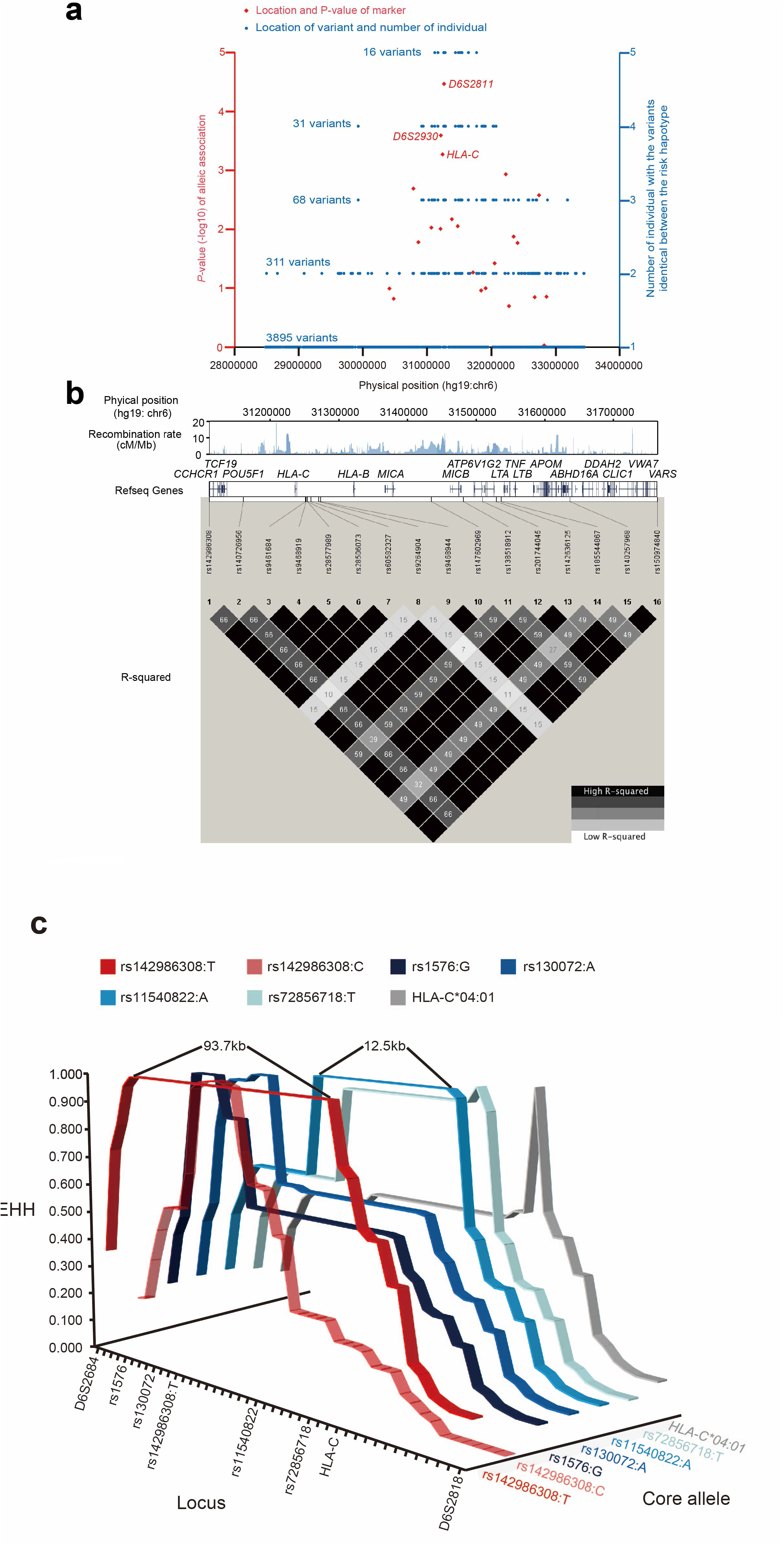
Identification of AA susceptibility variant by association and sequencing analysis. **a** Association analysis and risk haplotype resequencing of the MHC region. Red diamonds indicate P-values (-log10 scale) and locations. Three diamonds refer to haplotypes used for downstream analysis. Blue circles indicate individuals with variants identical between risk haplotype cases and the variant locations. **b** Pair-wise LD between 16 variants identified by NGS using genotype data from 89 Japanese individuals obtained with the 1000 Genomes Browser. Upper track shows recombination rate (cM/Mb) estimated from Phase II HapMap data (release 21), middle track the gene map (RefSeq genes) generated with the UCSC Genome Browser, and lower track the pair-wise LD between the 16 variants in R-squared. **c** EHH analysis of core alleles at 43 loci displaying LD with the T allele of rs142986308. An estimated 43 loci haplotypes encompassing 24 SNVs of CCHCR1 and 19 multi-allelic loci (2.32Mbp) were used for this investigation. The 7 selected core alleles were as follows: rs142986308 allele T, rs142986308 allele C for the internal control, 4 SNVs that displayed LD with rs142986308 (**Additional file1: Figure S14**), and *HLA-C*04:01* (**Additional file1: Figure S15**) as functional variants.

**Table 1.**
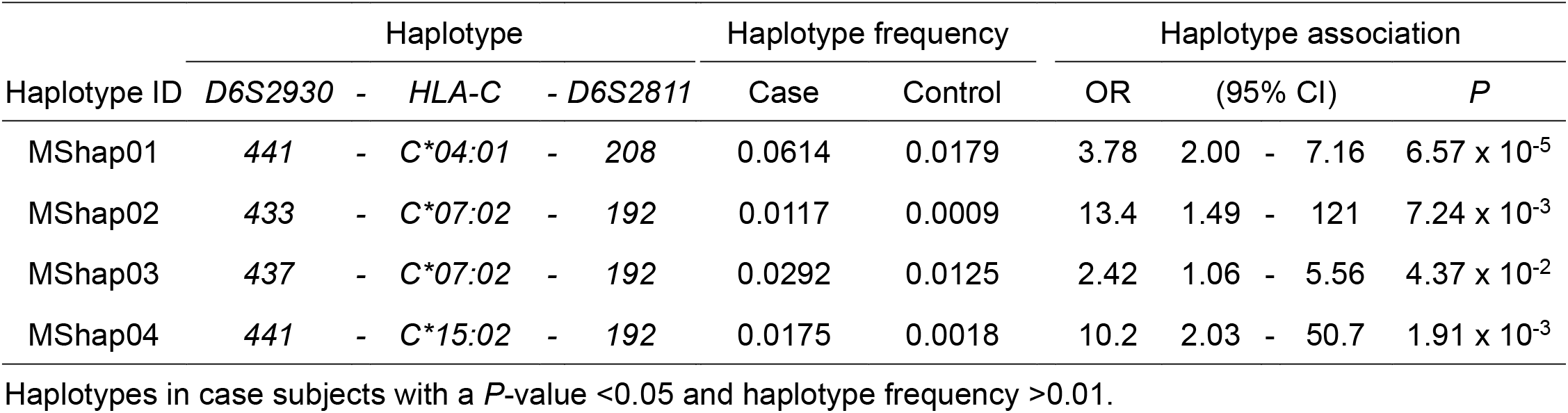
Haplotype association analysis of 3 loci around *HLA-C* gene.

To move from the identified risk haplotypes to causal AA variants, we next sequenced 5 individuals with MShap01 and 7 individuals with the other non-risk haplotypes spanning the entire MHC (chr6:28477797-33451433, hg19) (**Additional file1: Table S3**). As all risk haplotypes were heterozygous, any AA causal variant(s) would be expected to be heterozygous as well. Therefore, variants were accordingly filtered and only variants found to be identical between risk haplotypes were retained. Following this strategy, we extracted 3895 heterozygous risk variants from the 77,040 variants identified in the 12 individuals (**Additional file1: Table S4**). Of these, only 16 variants were identical between the 5 AA risk haplotypes (Fig. 1a, **Additional file1: Figure S2, and Table S5 and S6**) and only one was a non-synonymous coding SNV (rs142986308), defining a p.Arg587Trp substitution and mapping to *CCHCR1* (**Additional file1: Figure S3**). Using pair-wise LD analysis, we further established that a haplotype composed of 16 extracted variants in a 651.7-kb region between *CCHCR1* and *VARS* displayed strong LD despite including segments with a high recombination rate (Fig. 1b), implying that the haplotype was likely to be younger and/or has undergone positive selective pressure[16]. These results also suggested that our strategy used for stratification, sequencing, and filtering was effective for discovering risk haplotypes and novel MHC variants associated with AA.

### Sequencing *CCHCR1* gene and haplotype analysis

To check for further variations in *CCHCR1*, we sequenced all coding exons in all subjects and found 2 additional nonsense and 22 non-synonymous variants, though only SNV rs142986308 was shared between all 5 patients and demonstrated a significant association with AA (OR = 3.41, CI 95% = 1.94-5.99, *P* = 3.39 × 10^−5^) (**Additional file1: Table S7**). These two nonsense variants were mapped to the coding regions in alternative transcripts 1 and 2, which are common in Japanese and European Americans, and to the non-coding region in alternative transcript 3 (**Additional file1: Figure S4 and Table S7**). SNV rs3130453 gave rise to the shortest of the 3 alternative transcripts previously reported for *CCHCR1* [19] and SNV rs72856718, which has not been reported and is predicted to result in the same alternative transcript (**Additional file1: Figure S4**).

To confirm the T allele of variant rs142986308 as an AA susceptibility allele, we estimated haplotypes for 24 SNVs. Haplotype 26 (Hap26) harboring the T allele rs142986308 showed a statistically significant association with AA (OR = 3.41, CI 95% = 1.94-5.99, *P* = 3.39 × 10^−5^), and rs142986308 was the only SNV associated with AA in Hap26 (Table 2). Thus, rs142986308 was determined to be the primary variant associated with AA.

**Table 2.**
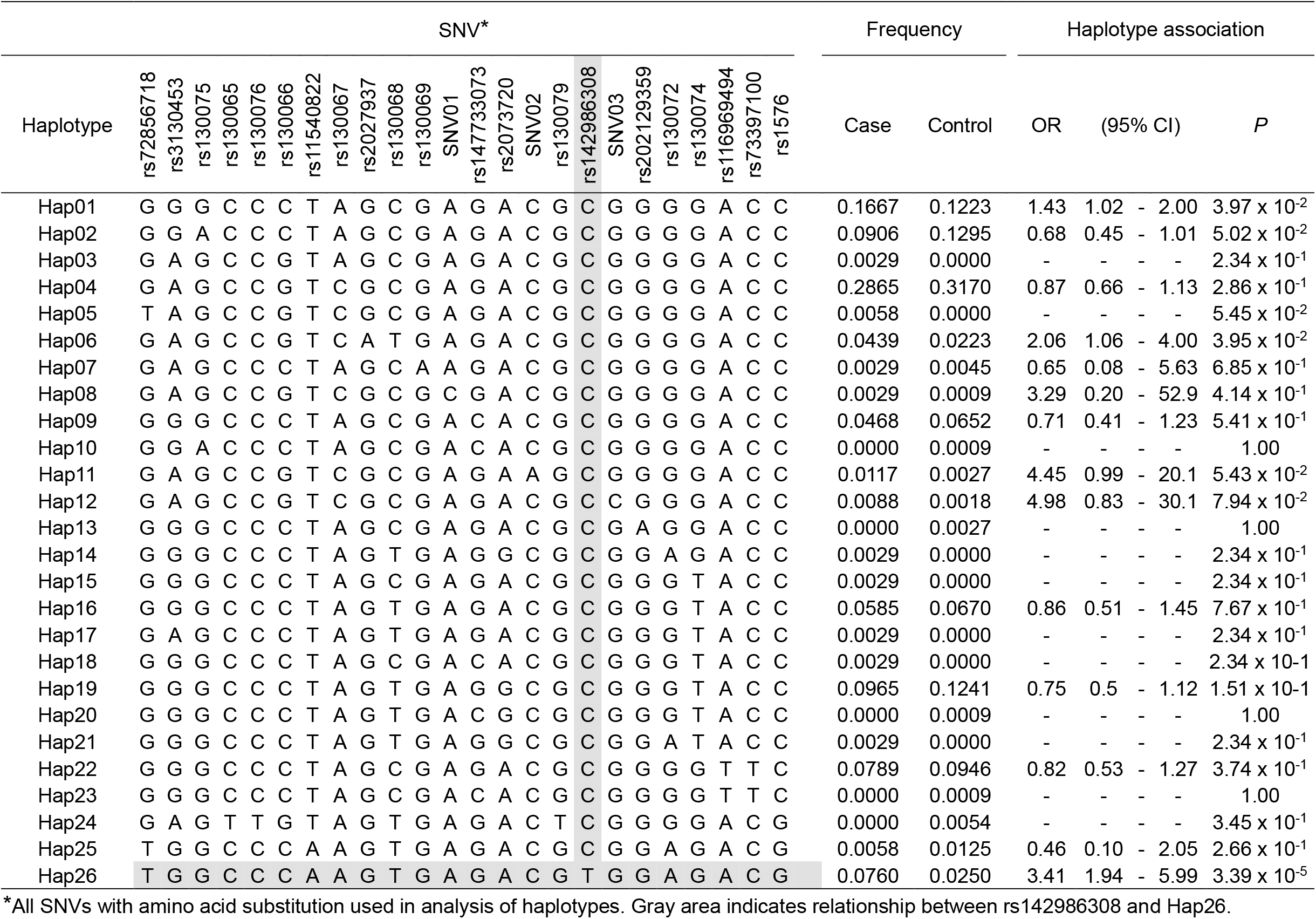
Haplotype analysis of 24 SNVs with amino acid substitution in *CCHCR1*.

Next, we investigated decay of the risk haplotype by recombination events that have occurred in evolutionary history. Extended haplotype homozygosity (EHH)[20] analysis with 5 of the identified risk alleles (designated as core alleles in Fig. 1c) was conducted. The core allele T of rs142986308 tagged the largest (93.7 kb) LD block with all values between rs1576 and *D6S2931* showing EHH=1.00 (Fig. 1 c), indicating that Hap26, exclusively shared by the patients, had recently increased in frequency. Adjacent alleles also showed some frequency increase in the patient group, possibly by hitchhiking (**Additional file1: Table S7**). Although the exact selection mechanism operating for the T allele of rs142986308 remains unknown and the overall number of haplotypes analyzed is modest, it is plausible that this allele is the primary target of selection for AA. Hence, we excluded the other nonsense and non-synonymous variants as causal for AA, despite being in strong LD with the causal rs142986308 variant identified by haplotype and EHH analyses.

### Evaluation of impact on function of CCHCR1 by allele T of rs142986308

We next examined whether the variant rs142986308 had influence on the function of CCHCR1, which is predicted to contain several coiled-coil domains (Fig. 2a **and Additional file1: Table S7**) [21]. The domain including the AA-associated variant p.Arg587Trp is well conserved in parts across many species (Fig. 2a **and Additional file1: Table S8**). Using structure prediction, the probability of coiled-coil conformation of this domain was notably reduced in only Hap26 harboring the AA-associated T allele of rs142986308 (Fig. 2b **and Additional file1: Figure S5 and S6**). Aromatic substitutions are known to be more disruptive towards coiled-coil domains than alanine, glutamic acid, lysine, leucine, and arginine, which favor coiled-coil domain formation[22], adding weight to our speculation that the variant rs142986308, which substitutes arginine with aromatic tryptophan, does indeed impair coiled-coil conformation of CCHCR1 (Fig. 2a).

**Fig. 2.**
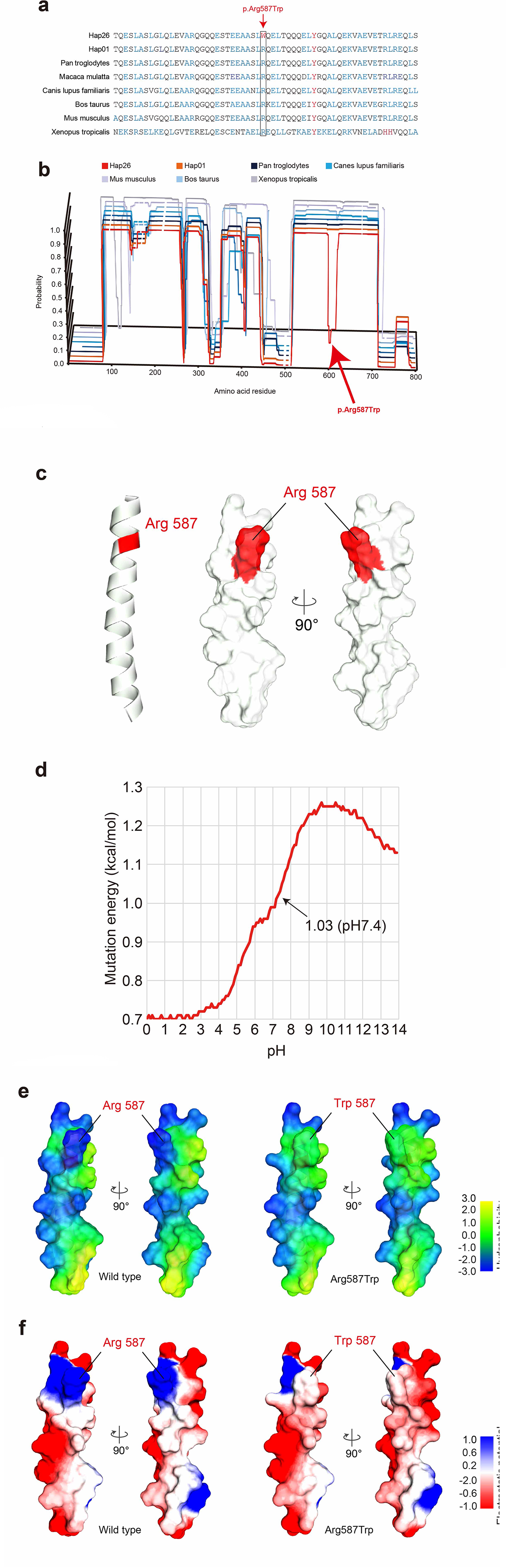
Prediction of impact on function of CCHCR1 by amino acid substitution. **a** Multiple amino acid sequence alignment of CCHCR1 showing evolutionarily conserved amino acids. The sequences, except for Hap01 and Hap26, were NP_001009009 (Pan troglodytes), NP_001108422 (Macaca mulatta), XP_532064 (Canis lupus familiaris), NP_001019707 (Bos taurus), NP_666360 (Mus musculus), and NP_001116918 (Xenopus tropicalis). Blue indicates residues that prefer to form coiled-coil domains (Ala, Glu, Lys, Leu, Arg) and red indicates aromatic residues that do not prefer to form coiled-coil domains. Arrow and box indicate the position of substitution of p.Arg587Trp (rs142986308). **b** Coiled-coil structure prediction of CCHCR1 in the AA-associated haplotype and different species using COILS v2.2. The Y axis indicates the probability of coiled-coil conformation and the X axis amino acid residue number. Full amino acid sequences are described in **Additional file1: Supplementary sequences**. Multiple amino acid sequence alignments were assigned to the probabilities of coiled-coil conformation for each haplotype and specie. Line brakes correspond to gaps in the multiple alignments. Arrow shows position of p.Arg587Trp (rs142986308). **c** Ribbon and molecular surface representations of partial CCHCR1 structure. Surface models are shown in 2 orientations. Residue 587 is represented in red. **d** Effects of Arg587Trp mutation on CCHCR1 protein structure at different pH values. A change of >0.5 cal/mol, which destabilized the structure, was classified as a significant change. Surface maps showing **e** hydrophobicity and **f** electrostatic potential of the wild type and Arg587Trp are presented.

We also performed homology searches using the PDB database and obtained the template structure for partial CCHCR1 protein around the p.Arg587Trp substitution, the crystal structure of the human lamin-B1 coil 2 segment (Fig. 2c) [23]. We then generated a partial CCHCR1 structure using the template and evaluated the effect of p.Arg587Trp substitution on protein stability by performing molecular dynamics simulations using a CHARMm force field. The simulation showed that p.Arg587Trp substitution reduced protein stability (mutation energy: 1.03 kcal/mol) (Fig. 2d) and changed the CCHCR1 structure around the residue at 587 (Fig. 2e and f). This indicates that p.Arg587Trp substitution may alter the protein-protein interaction of CCHCR1 (Fig. 2c).

### Allele-specific genome editing using CRISPR/Cas9 in mice

Next, we functionally evaluated the AA-associated variant by *in vivo* phenocopying p.Arg591Trp in murine *CCHCR1* using allele-specific genome editing with the CRISPR/Cas9 system. Mice were generated with the risk allele concordant with p.Arg587Trp derived from the T allele of rs142986308 in humans (Fig. 3), then we established mouse strains with homozygous risk alleles (AA mice). In this study, 15 mice (55.5%) among 27 AA mice displayed patched dorsal hair loss until10 months after birth, thus successfully phenocopying the AA phenotype. The incidence of hair loss was higher in those mice as compared to C3H/HeJ mice, which show spontaneously development of AA with age. [24] Over time, the initial area of hair loss expanded in the majority of the AA mice (Fig. 4a), though constant and recovered hair loss was observed in some of those mice (**Additional file1: Figure S7 and S8**). The male to female ratio of alopecic AA mice was nearly equal, and their surface displayed black spots, while the hairs appeared to be broken and tapering (Fig. 4b), similar to the conditions seen in humans with AA and specific for AA-associated hair loss[25]. Thus, an altered hair shaft structure may be a feature concordant between human AA[26] and the present engineered AA mice. All of the AA mice showed retained hair follicles in the area of hair loss and no signs of lymphocyte infiltration were seen in microscopic observations (Fig. 4c), implying involvement of a non-immune mechanism.

**Fig. 3.**
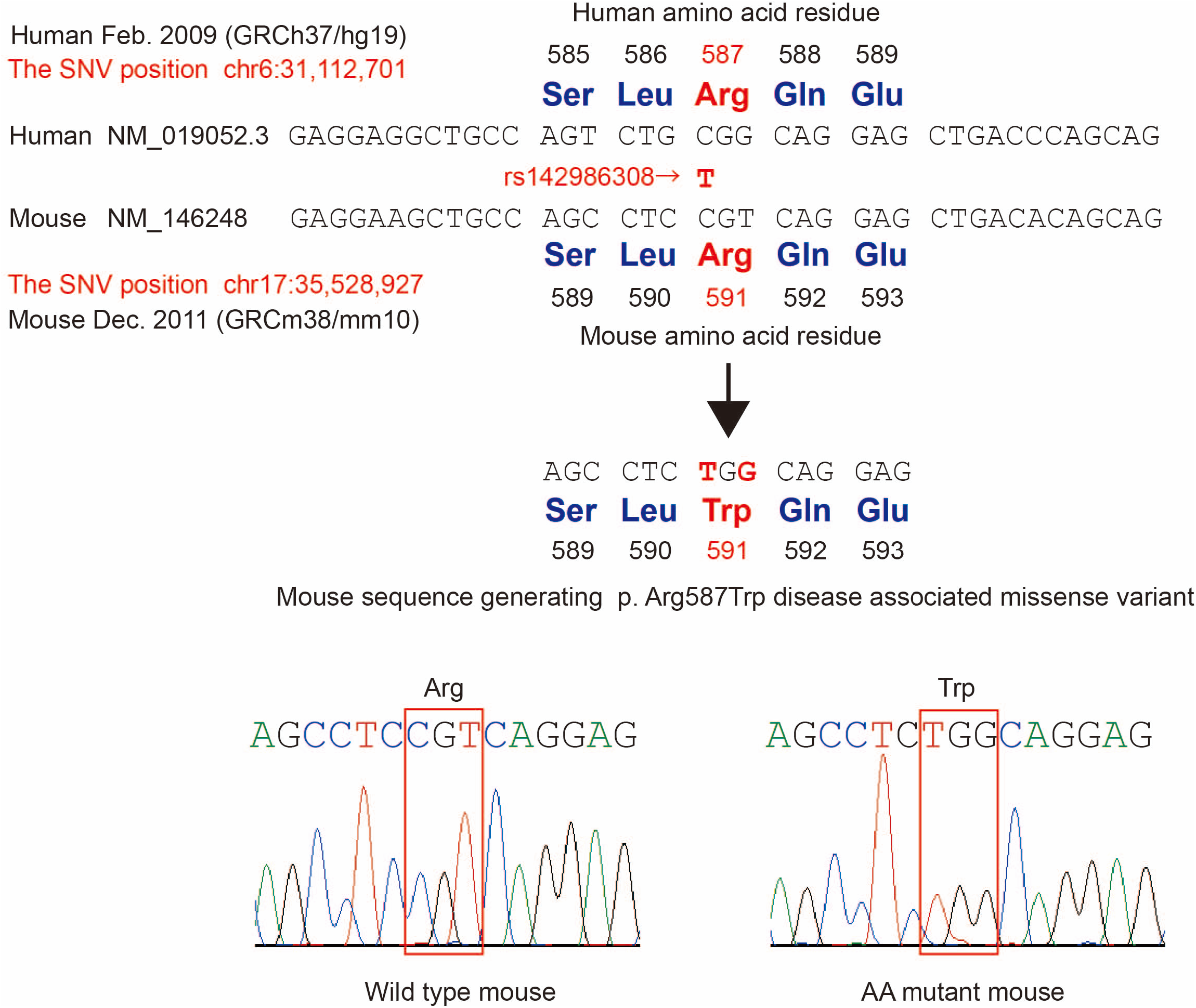
Sequence alignments noted in human and mouse samples around AA-susceptibility variant. Upper portion shows the mouse sequence concordant for Arg587 amino acid in human and mouse sequences aimed for allele-specific genome editing. Lower portion shows electropherograms from sequencing for genotyping the allele.

**Fig. 4.**
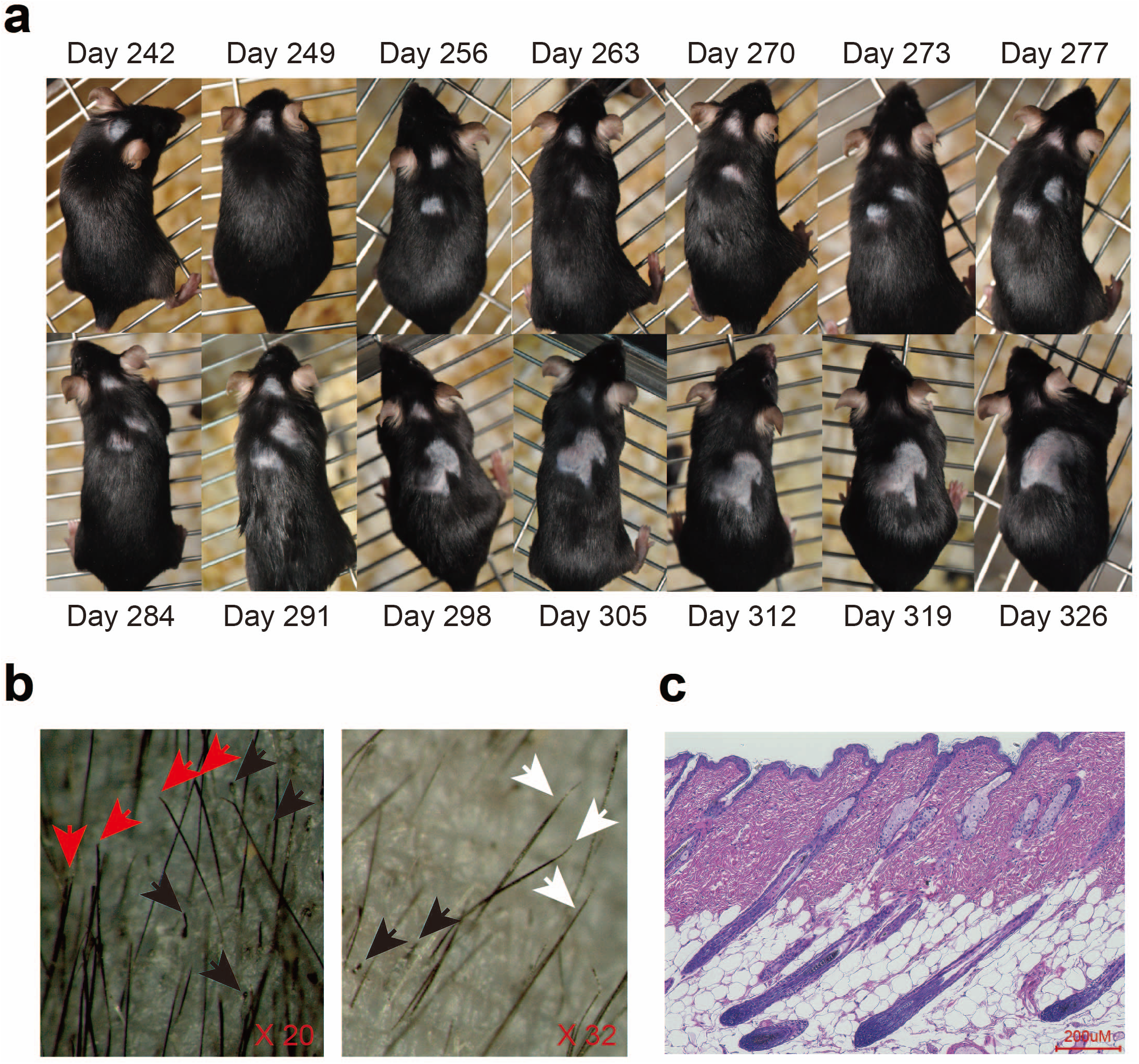
Phenotypic traits of skin biopsy samples from AA risk allele mice (AA model mice). **a** Expansion of hair loss area in representative AA mouse. **b** Morphology of hair loss area in AA mouse. Red arrows show broken hair, black arrows show black spots, white arrows show tapering hair. **c** Microscopic features of hair loss area in representative AA mouse. Paraffin section of skin from representative AA mouse after staining with hematoxylin and eosin.

### Microarray analysis of alopecic mice skin biopsies

To investigate the presence of a possible biological function underlying the observed hair loss in the AA mice, we performed gene expression microarray analysis of dorsal and ventral skin biopsies (**Additional file1: Figure S9 and S10**), which resulted in identification of 265 probes (246 genes) with 2-fold or greater up- or down-regulation as compared to wild-type mice. The full list of genes is shown Table S11. Clustering analysis of these probes uncovered a strongly up-regulated gene cluster in the AA mice (Fig. 5a), including hair-related genes (Fig. 5b). Nearly all of those up-regulated with a greater than 25-fold change value were keratin[27] (n=12) and KRTAP (n=31) genes (**Additional file1: Table S9**). Hair keratins and KRTAPs are the major structural components of the hair shaft, and specifically expressed in the medulla, cortex, and cuticle layers of the shaft[28], and interaction between hair keratins and KRTAPs suggest their contribution to its rigidity[29].

**Fig. 5.**
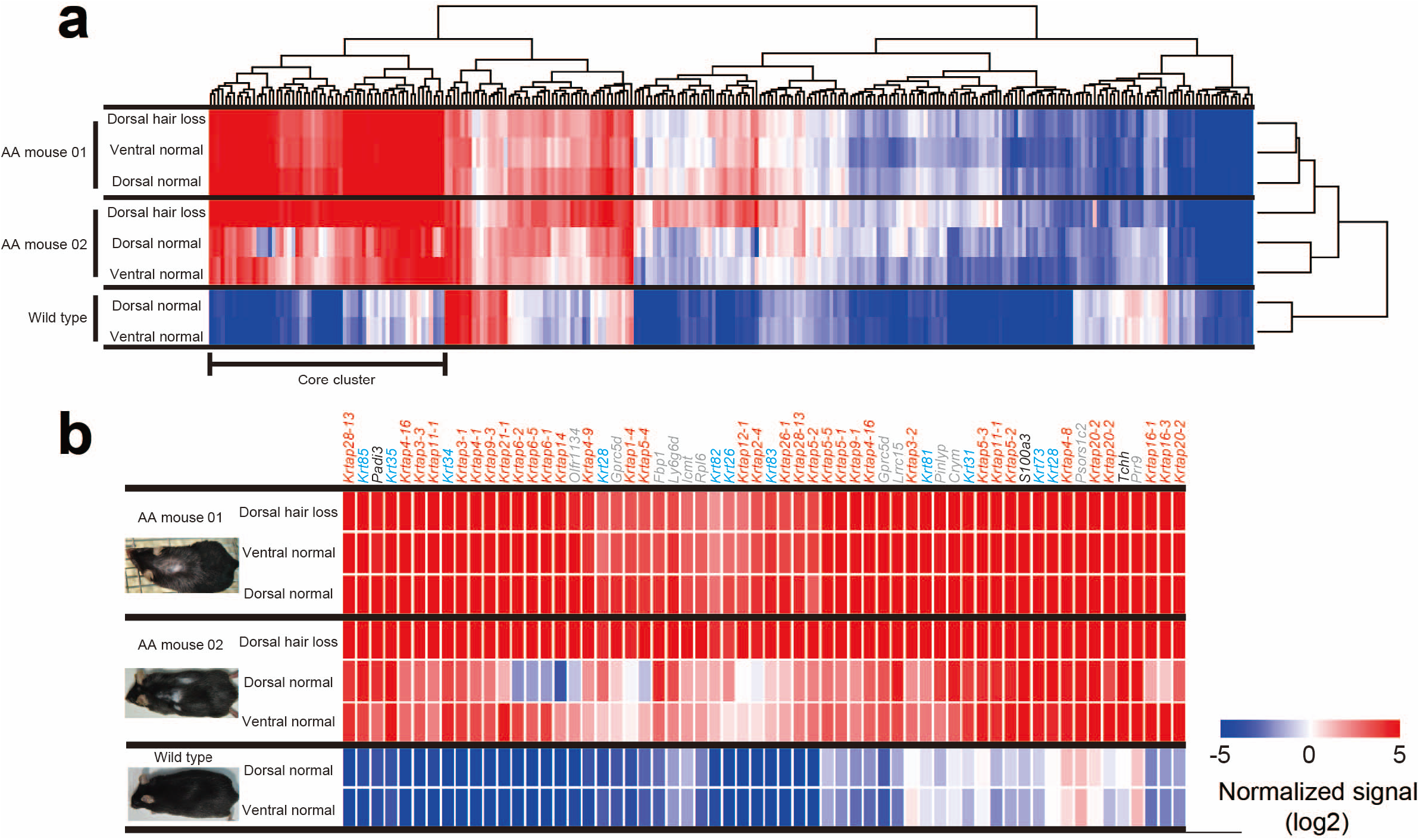
Microarray analysis of skin biopsy samples from hair loss areas in 2 AA mice and 1 wild-type mouse. **a** Heat map of 265 probes showing ≥2-fold change in gene expression. The list of genes is shown in Table S11. The cluster displaying high expression in the dorsal hair loss area in both AA mice was defined as the ‘core cluster’. **b** Heat map of core cluster genes. The color code depicts KRTAP family (red), keratin family (blue), other hair-related (black), and non hair-related (grey) genes.

Other up-regulated genes included peptidyl arginine deaminase type III (*Padi3*), S100 calcium binding protein A3 (*S100A3*), trichohyalin (*Tchh*)[30], and homeobox C13 (*Hoxc13*)[31] (**Additional file1: Table S9**). In cuticular cells, S100A3 is a substrate of PADI3, while in inner root sheath (IRS) cells of hair follicles it is a substrate of TCHH[30]. Ca^2+^-dependent modifications of S100A3 and TCHH by PADI3 play important roles in shaping and mechanically strengthening hair with hair keratin[30, 32]. *Hoxc13* is unique among the Hox genes, as it is expressed in the outer root sheath (ORS), matrix, medulla, and IRS of hair follicles in a hair cycle-dependent manner, and it has a role in hair shaft differentiation[31]. Thus, the majority of genes shown to be strongly up-regulated in AA mice are involved in the hair shaft and its formation.

Hairless model mice, nude mice lacking forkhead box N1 (*Foxn1*), have a pleiotropic mutation that leads to 2 independent phenotypic effects, which are disturbed development of hair follicles and dysgenesis of the thymus[33]. The *Foxn1* expression has been shown to be regulated by *Hoxc13* in hair follicles[34], and both *Hoxc13-null* and *Hoxc13*-overexpressing transgenic mice were found to be an alopecic phenotype[34]. These three alopecic mice lines display defective hair shafts or aberrant hair cuticles[33, 35, 36]. In the present microarray analysis, *Foxn1* expression in alopecic skin of AA mice was also upregulated by 6.5-fold as compared to the wild-type mice, though this gene was ruled out by the filtering criteria employed (**Additional file1: Figure S11**). A comparison between gene profiling results of *Hoxc13*-null[34] and AA mice indicated that the regulated genes were quite similar between them (Fig. 6 and **Additional file1: Table S10**). Thus, among 113 down-regulated genes with a gene symbol in *Hoxc13*-null mice[34], 89 were upregulated in the AA mice (**Additional file1: Table S10**), the majority of which were hair keratins or KRTAPs. Moreover, we observed a significant inverse correlation of fold change values for these genes between these mouse lines (Fig. 6), despite differences in strain and mutant genes. Therefore, *Cchcr1* may be involved in the hair shaft differentiation regulatory network controlled by *Hoxc13, Foxn1*, keratins, and KRTAPs, though the regulation was in opposite directions.

**Fig. 6.**
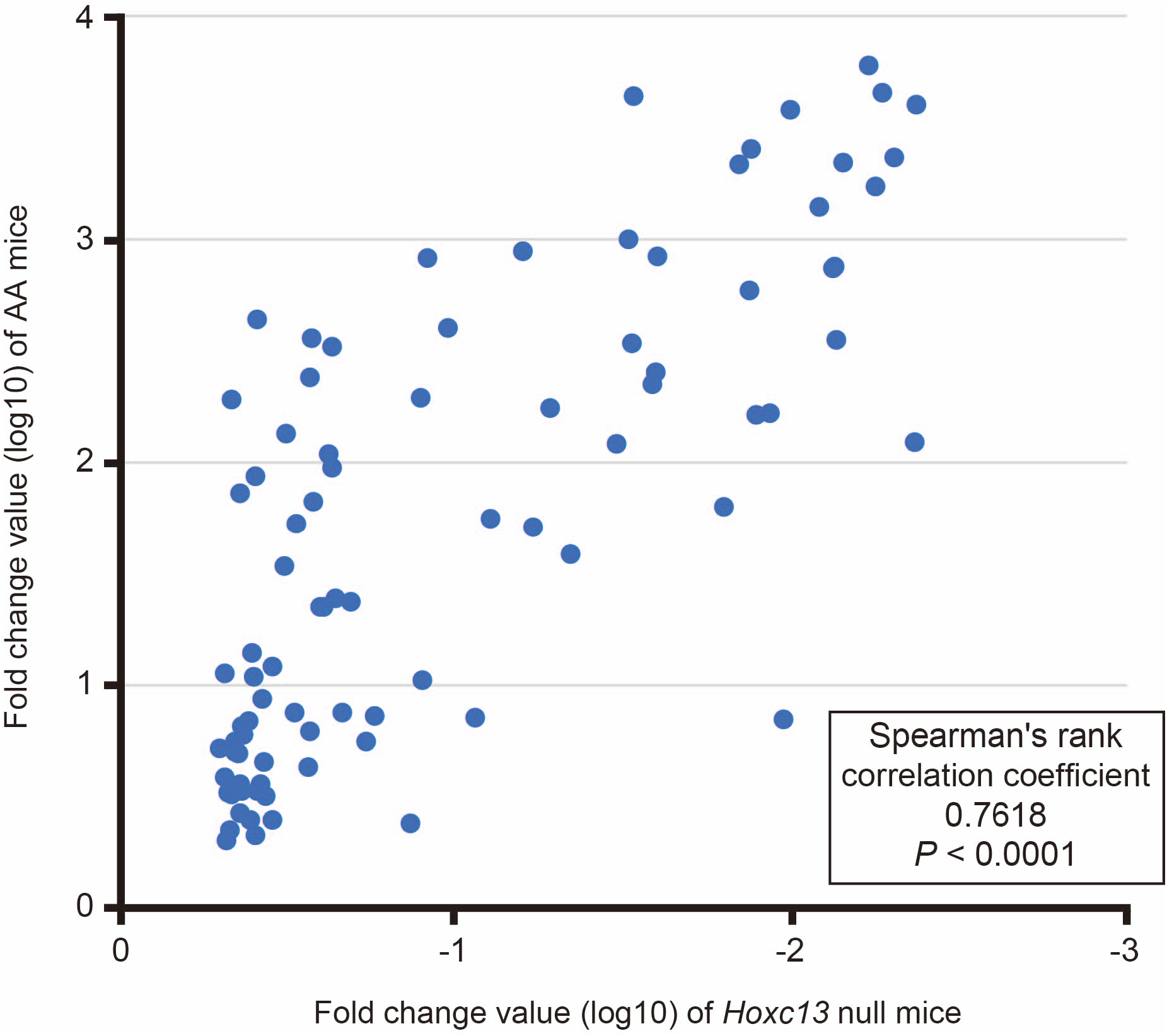
Absolute fold change values of 89 concordant genes showing ≥2-fold change in *Hoxc13* null and AA mice as compared to wild type. All genes demonstrated down-regulation in *Hoxc13* null mice and up-regulation in AA mice. Correlation coefficients are indicated for each mouse strain. Correlation coefficients and statistical significance in comparisons between *Hoxc13* null and AA mice were determined using Spearman’s rho.

Enrichment analysis of these 246 genes using the Database for Annotation, Visualization and Integrated Discovery (DAVID) tool[37] showed Gene Ontology[38] (GO) terms relating to only keratin and the intermediate filament, but not to any immunological pathways (**Additional file1: Table S11**).

To confirm the microarray results, we performed quantitative PCR (qPCR) analysis of 7 of the up-regulated genes and *Cchcr1* using a *comparative C_T_* method[39]. Those results confirmed significant differences between dorsal skin biopsies from areas of hair loss in the AA and wild-type mice (Fig. 7a **and Additional file1: Figure S12**). All AA mice showed highly concordant rank orders of expression levels of the examined genes (Fig. 7b **and Additional file1: Figure S13**), implicating involvement of the regulatory network in hair shaft differentiation[34].

**Fig. 7.**
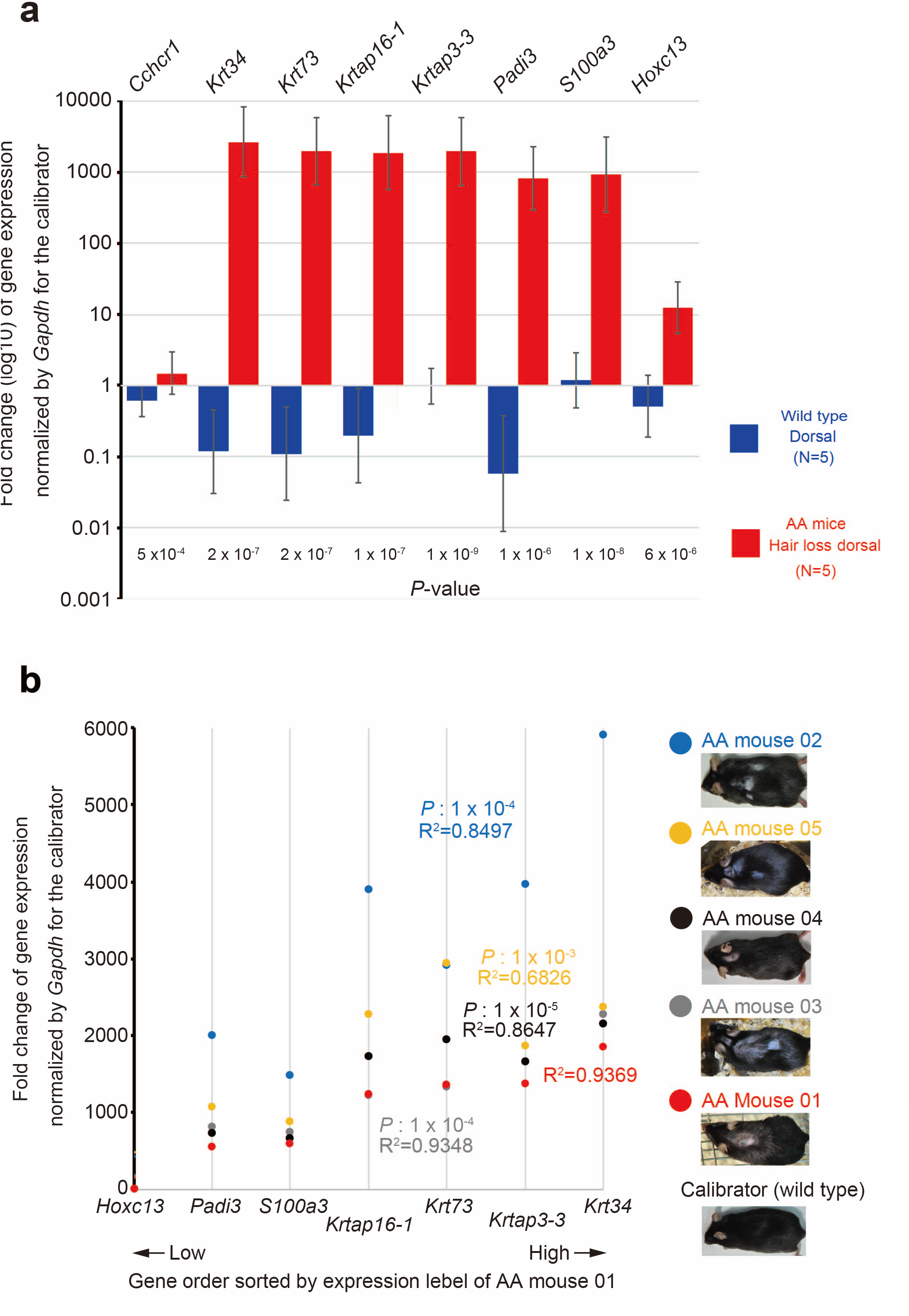
Validation of upregulated gene expression for Cchcr1 and 7 selected genes. **a** Mouse skin biopsy samples were subjected to expression analysis by qPCR and a comparative CT method. Bars reflect 95% confidence intervals. Fold change values were normalized to dorsal hair loss in a wild-type mouse as a calibrator, thus the fold change value of the calibrator was always 1. Statistical significance was determined using Student’s t test. **b** Gene expression trends in skin biopsy samples from dorsal hair loss areas in AA mice. Correlation coefficients are indicated for each mouse. Statistical significance was determined using Pearson’s product-moment correlation between each mouse and AA mouse 01.

### Hair shaft and follicles

Finally, we evaluated the localization of CCHCR1 using immunostaining and electron microscopy to delineate potential mechanisms underlying the observed hair loss in our mouse model. Immunostaining revealed CCHCR1 to be located in the mid-to-upper hair shaft but not in the cell and hair shaft within hair bulb (Fig. 8a-i), suggesting that CCHCR1 is a structural component, similar to keratin, of the hair shaft. CCHCR1 was co-localized with the hair cortex and also found to be localized in the hair medulla (Fig. 8b, e and h). Expression of this protein in the skin of AA mice was stronger as compared to that of the wild type (Fig. 8a-i), supporting our findings of expression analysis using RNAs (Fig. 5a, b). Hair keratins are known to be organized in bundles with structures dominated by alfa-helical coiled-coils[40]. Using low-resolution immunostaining, we were not able to identify any difference between the AA and wild-type mice. However, high-resolution scanning electron microscopy (SEM) results demonstrated hair abnormalities, including turbulence in cuticle formation and flattened shafts in alopecic AA mice (Fig. 8p-x). Alopecic AA mice displayed aberrant hair not only in areas of hair loss (Fig. 8×-t), but also in normal areas (Fig. 5s-u). Moreover, AA mice that did not yet show hair loss also displayed aberrant hair (Fig. 8p-r), suggesting that all AA mice are affected by abnormal hair keratinization. These aberrations are similar to hair cuticle aberrations observed in AA patients[41], as well as in *Foxn1*-null, *Hoxc13*-null, and *Hoxc13*-overexpression mice[33, 35, 36].

**Fig. 8.**
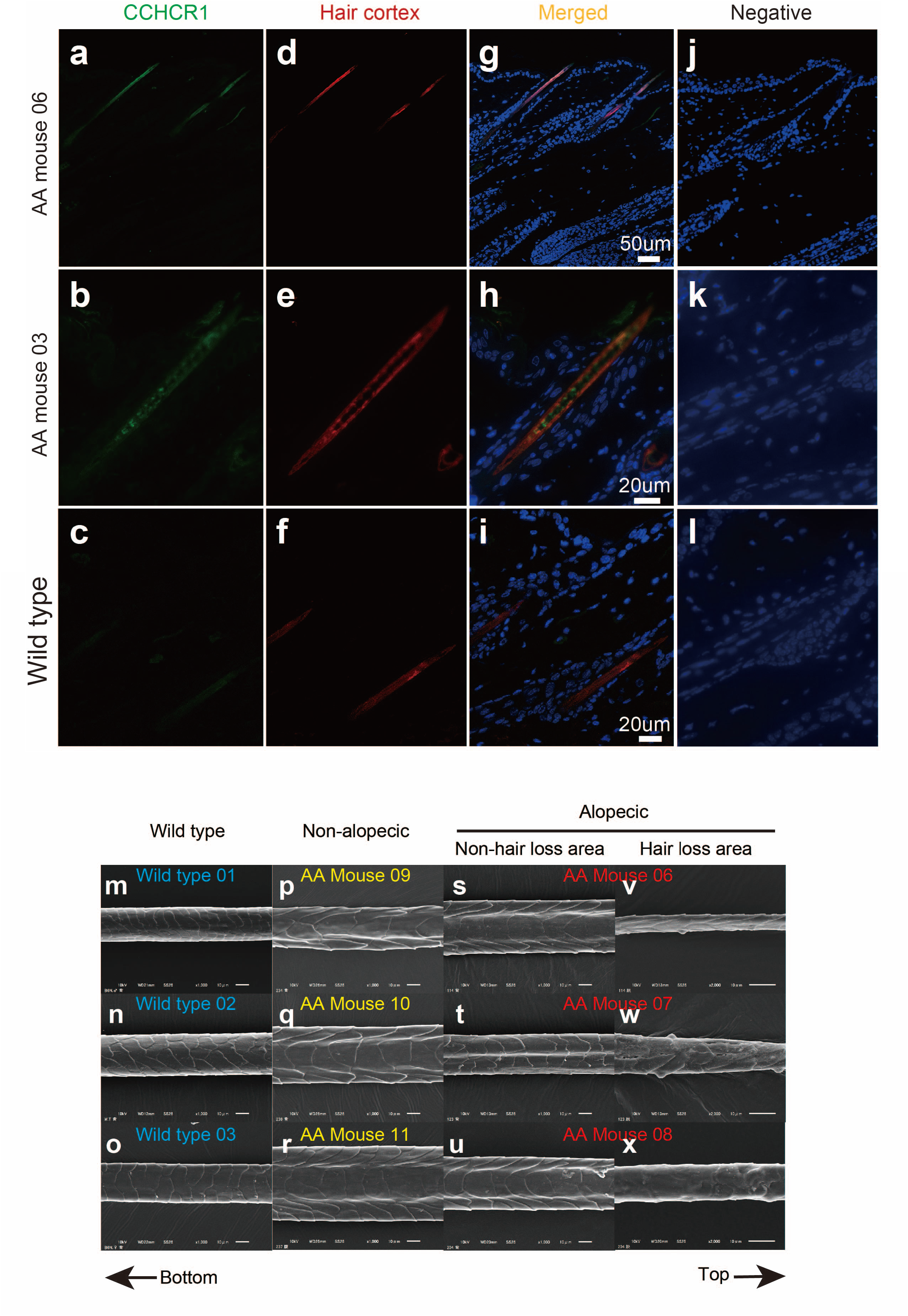
Investigations of hair shafts and follicles. **a-l** Co-localization of CCHCR1 with hair cortex keratin in follicles from AA and wild-type mouse skin. Paraffin sections were stained with anti-CCHCR1 and anti-hair cortex keratin antibodies, and subjected to fluorescent microscopy. Left panels (**a, e, i**) show CCHCR1 (green). Middle panels (**e, g, k**) show hair cortex keratin (red). Right panels (**c, g, k**) show merged sections with DAPI nuclear staining (blue). Upper panels (**a-d**) show longitudinal section of skin including subcutaneous tissue from AA mouse. Middle panels (**e-h**) show longitudinal section of upper hair follicle from AA mouse. Lower panels (**i-l**) show longitudinal section of upper hair follicle from wild-type mouse. (**m-x**) Scanning electron microscopy (SEM) imaging of hair shafts on dorsal skin was performed. In each panel, hair orientation is shown with the bottom to the left and top to the right. Non-alopecic indicates AA mice that have not yet displayed hair loss (pr). Alopecic indicates AA mice that have already displayed hair loss (**s-x**). Alopecic column also indicates hairs from non-hair loss (**s-u**) and hair loss (**v-x**) areas in the same mouse. Data shown in 3 columns are from 3 different mice.

## Discussion

Our finding that *CCHCR* is an AA-associated locus in the HLA class I region differs from that of a previous genome-wide association study, which suggested that *HLA-DR* in the HLA class II region is a key driver of AA aetiology[13]. One possible explanation for this seeming contradiction is that our study may not have been sufficiently powered to detect the weak association with AA within the HLA-class II region, though the microsatellites utilized covered the entire class II region as well (Fig. 1a). On the other hand, consistent with our finding, studies performed in Chinese using HLA genes as markers suggested that the locus associated with AA maps to the class I rather than the class II region[42–44]. Moreover, variant rs142986308 is very rare in Caucasians (**Additional file1: Table S8**) and has undergone positive selection (Fig. 1c), at least in Japanese, suggesting population-specific differences regarding AA risk haplotypes. This is the first study to show that an HLA class I allele, rs142986308, can be functionally linked to the AA phenotype, which has not been demonstrated for any other variant, including *HLA-DR*.

Previous studies have provided evidence of a relationship between *CCHCR1* and hair keratin-related genes. For example, a risk haplotype (*CCHCR1*WWCC*) was previously implicated to be involved in psoriasis in Europeans[21, 45]. Transgenic mice with the risk haplotype appeared normal, though it was shown that overexpression of CCHCR1 affected keratinocyte proliferation[46], and hyperproliferation of keratinocytes is a hallmark for psoriasis. However, keratin-related genes showed altered expression in mice at risk and most of the genes with significantly lower expression were those encoding hair keratins or KRTAPs[47]. These observations suggested that expression of these genes is dependent on the *CCHCR1* haplotype.

AA shows clinical heterogeneity[48], thus findings from the lymphocyte infiltration model as well as the AA mice are not able to fully explain the pathogenesis. Those obtained with AA mice may be more suitable to explain aberrant hair shaft results, such as black dots (cadaverized hairs), tapering hairs, and broken hairs, which are specific AA-associated hair loss conditions[25]. Therefore, investigation of the relationship between autoimmunity response and aberrant keratinization in hair follicles may lead to a better understanding of AA pathogenesis. Conversely, each model may have an independent pathway, leading to AA heterogeneity.

The current study has several limitations thus further studies are needed. First, more detailed investigation of immunological function in AA mice must be performed, because the present study did not completely exclude immunological aspects. Second, additional examination of factors that eventually lead to hair loss is needed, though all of the AA mice displayed abnormal hair shafts. In addition, investigation of how the mutation alters the function of the CCHCR1 protein is necessary. Finally, it is important to investigate hairs and hair follicles obtained from patients with and without the risk allele in order to fully understand the functions of CCHCR1 in human hair.

In summary, we identified a novel AA susceptibility variant (T allele of rs142986308, p.Arg587Trp) in the human MHC class I region and established an alopecic mouse model. Findings of functional analyses indicate that CCHCR1 is a novel component in hair shafts and its variation alters the coiled-coil formation of CCHCR1, resulting in abnormalities of hair shafts and cuticles, along with up-regulation of hair keratin, KRTAPs, and other relevant genes, ultimately leading to AA pathogenesis, shown by fragile and impaired hair as well as hair loss. In addition, our findings suggest an alternative pathway for that pathogenesis based on aberrant keratinization. Together with the present engineered mice, our results provide a valuable resource for continued research of AA pathogenesis and development of potential future treatments.

## Conclusions

This study genetically identifies an AA susceptibility variant by association and sequencing analysis within the MHC region, and functionally confirms that the variant involves hair abnormalities using engineered mice by CRISPR/Cas9 for allele-specific genome editing.

## Materials and methods

### Patients and controls

Upon approval of the experimental procedures from the relevant ethical committees of Juntendo University and Tokai University, we obtained informed consent from all unrelated AA and healthy individuals prior to collection of DNA samples. A total of 171 individuals affected with AA (MAA-multiple alopecia areata: 115, AT-alopecia totalis: 18, AU-alopecia universalis: 38) and 560 unrelated individuals of Japanese origin participated in this study. All cases were diagnosed and treated at a Juntendo University Hospital in Japan. DNA was extracted using a QIAamp DNA blood kit (QIAGEN, Hilden, Germany) under standardized conditions to prevent variations in DNA quality. For additional quality control, we used 0.8% agarose gel electrophoresis to check for DNA degradation and/or RNA contamination, and performed optical density measurements to check for protein contamination. The final DNA concentration was determined with 3 successive measurements using a PicoGreen fluorescence assay (Molecular Probes, Thermo Fisher Scientific, Rockford, IL, USA).

### Microsatellite genotyping

We selected 22 microsatellites spanning 2.44 Mbp (from *HLA-E* to *PSMB9* gene) in the HLA region (**Additional file1: Table S12**) harbouring the HLA class I, II, class III regions, and genotyped all patients and control subjects. Forward primers of the primer sets to amplify microsatellites were labeled by 5’ fluorescent FAM. Oligonucleotides were obtained from Greiner Bio-one. PCR and fragment analyses were performed with capillary electrophoresis using an Applied Biosystems 3730 Genetic Analyzer. Allele assignment was determined with GeneMapper Software (Thermo Fisher Scientific) and conducted as previously described[49]. Fragment sizes were assigned to allele names in the corresponding microsatellites. In the *MICA* locus, 5 *MICA* polymorphisms (*A4, A5, A5.1, A6, A9*) were determined based on the number of alanine (GCT) repeats. The A5.1 allele contained 5 GCT repeats, plus 1 extra guanine nucleotide (GCT)_2_G(GCT)_3_.

### *HLA-C* locus genotyping

A LABType^®^ SSO typing test produced by ONE LAMBDA (Inc., Canoga Park, CA) was utilized. This product is based on the reverse SSO method for use with a suspension array platform with microspheres as a solid support to immobilize oligonucleotide probes. Target DNA is amplified by PCR, then hybridized to the bead probe array, followed by flow analysis using a LABScan^TM^ 100 flow analyzer (ONE LAMBDA). *HLA-C* locus genotype data from 156 AA patients and 560 controls obtained in our previous study[18] were used and we also genotyped an additional 15 AA cases as part of the present study.

### Genomic library construction and sequencing

For HLA region capture and sequencing, genomic DNA (2 μg) was sheared to approximately 500 bp in size using a Covaris Acoustic Adaptor. Genomic libraries were prepared using a TruSeq DNA Sample Preparation kit. v.2 (Illumina, San Diego, CA, USA) following the manufacturer’s instructions, which involved size selection of DNA fragments of 550-650 bp in length on 2% agarose gels. HLA region enrichment was performed with an adaptor-ligated DNA sample library using the SeqCap EZ Choice Library Human MHC Design system (Roche NimbleGen, Madison, WI, USA) [50], according to the manufacturer’s instructions. To quantify and verify the genomic libraries, all samples were analyzed with a Bioanalyzer 2100 (Agilent Technologies, Santa Clara, CA, USA) using an Agilent DNA 1000 kit prior to sequencing (**Additional file1: Figure S16**). Sequence analysis was performed with the Illumina Genome Analyzer IIx platform, using a paired-end sequencing protocol (2×100 bp). We sequenced 5 risk and 7 non-risk haplotypes to a mean depth of 249 reads (**Additional file1: Table S3**), covering 95.2% (mean) of the 4.97-Mb MHC region (chr6:28477797-33451433, hg19) with at least 10 reads (**Additional file1: Table S3**).

### Next generation sequencing (NGS) data analysis

Fastx-toolkit, v. 0.0.13 (http://hannonlab.cshl.edu/fastx_toolkit/index.html), was used for quality control of the sequencing reads. Reads that passed quality control were mapped to the human reference genome (UCSC Genome Browser assembly GRCh37/hg19, http://genome.ucsc.edu/) using Burrows-Wheeler Aligner (BWA), v. 0.5.9, with the default parameters[51]. After alignment, Sequence Alignment/Map (SAMtools), v. 0.1.17, was used to convert the .sam to .bam files[52], and potential PCR duplicates were flagged with Picard MarkDuplicates (v. 1.88; http://picard.sourceforge.net/). A Genome Analysis Toolkit (GATK, v. 2.2-8) was used to perform local realignment, map quality score recalibration, and variant detection[53]. SNVs and indels were then annotated for functional consequences at the gene and protein sequence levels using ANNOVAR[54]. Finally, we manually checked the raw sequencing data using Tablet, a sequence assembly visualization tool[55] (**Additional file1: Figure S2**).

### Variant discovery and genotyping of *CCHCR1* by Sanger sequencing

Coding exons of *CCHCR1* were sequenced using PCR-based capillary Sanger sequencing. Oligonucleotides were purchased from Greiner Bio-One (**Additional file1: Table S13**). PCR was performed in a reaction volume of 10 μl containing 5 ng of genomic DNA, 0.2 U of KOD FX Neo (TOYOBO Life Science, Osaka, Japan), 5 μl of 2× PCR Buffer, 2μ of dNTP (2 mM each), and 0.2 μM (final concentration) of each of the primers. The thermal cycling profile was as follows: initial denaturation at 94°C for 2 minutes and 35 rounds of amplification at 98°C for 10 seconds, then 59-65°C (depending on primer set; see **Additional file1: Table S13**) for 30 seconds and 68°C for 1 minute. PCR products were purified using an AMPure XP (Beckman Coulter, Fullerton, CA, USA), according to the manufacturer’s protocol. Purification and sequencing of the PCR products were carried out using a BigDye Terminator v3.1 Cycle Sequencing kit (Thermo Fisher Scientific) and BigDye XTerminator Purification Kit (Thermo Fisher Scientific), following the manufacturer’s instructions. Automated electrophoresis was performed with an ABI PRISM 3730 Genetic Analyzer (Thermo Fisher Scientific). Sequencing data were analyzed using Sequencher (v. 5.1, Gene Codes Corporation, Ann Arbor, MI, USA). Genomic coordinates for all variants were called using the UCSC Genome Browser assembly GRCh37/hg19 (http://genome.ucsc.edu/).

### Statistical analysis

Logistic regression models were used to assess the genetic effects of multi-allelic loci, SNVs, and AA risk haplotypes. Comparisons of genotype and haplotype frequency differences were done by regression analysis for log-additive models[56]. Unadjusted odds ratio (OR) and 95% confidence intervals (95% CI) were calculated. Analysis was carried out using the SNPassoc R library[56]. For these association analyses, we used Bonferroni-corrected values to account for the problem of multiple testing to a threshold P value of 1.98×10^−04^, after accounting for multiple testing of 252 alleles in 23 multi-allelic loci for the first microsatellite analysis. An exact P-value test of Hardy-Weinberg proportion and evaluation of LD (linkage disequilibrium) for multi-allelic loci were simulated by the Markov chain method within Genepop[57]. To evaluate the degree of LD for bi-allelic loci (SNVs) we used Haploview 4.2.[58] To estimate haplotypes for SNVs and multi-allelic loci, we used PHASE v2.1.1.[59] For EHH analysis, haplotype data were generated from 22 nonsynonymous SNVs, 2 stop gain SNVs, and 19 multi-allelic loci with fastPHASE v1.2.[60] Moreover, for this estimation, each multi-allelic locus was regarded as a SNV[15]. Thus, we were able to extract the allele demonstrating LD (D’ ≥0.5) for the AA risk allele (rs142986308: T allele) from all alleles in each locus, if such an allele was detected (**Additional file1: Figure S15**), and then merged it with the other alleles. EHH was calculated using the “rehh” package[61]. To estimate the power of this study design to detect associated loci with AA, we performed statistical power calculations using the Genetic Power Calculator web application[62] (http://pngu.mgh.harvard.edu/~purcell/gpc/) assuming a type I error of alpha (a) = 0.05. The statistical power for a significance level of a = 0.05 in our sample was calculated for the *D6S2930* locus under several assumptions, as follows: a) high risk allele frequency of 0.144 in the general population, b) prevalence for AA of 0.001, c) heterozygote genotype relative risk of 1.76, d) homozygote genotype relative risk of 4.66, and e) a control group of subjects comprised of unselected individuals from the general population not screened for AA. Based on these conditions, the statistical power for the *D6S2930* locus was calculated to be 0.781.

### Prediction of coiled-coil domain and structural analyses

Multiple sequence alignment was performed with CLUSTALW (http://www.genome.jp/). To predict coiled-coil domains within the amino acid sequence of CCHCR1, we used 2 different programs, COILS[63] (http://embnet.vital-it.ch/software/COILS_form.html) and Paircoil2[64, 65] (http://groups.csail.mit.edu/cb/paircoil2/), with the following recommended default settings: COILS v2.2 with a window size of 28 and the MTIDK table, and Paircoil2 with a window size of 28 and P-score cutoff of 0.025.

The amino acid sequence of CCHCR1 (GenBank accession: NP_061925.2) was obtained from the NCBI GenBank database (https://www.ncbi.nlm.nih.gov) and used as a target for homology modeling. After PSI-BLAST searches[66] of protein databank (PDB) sequence entries (http://www.rcsb.org/pdb/) using the CCHCR1 sequence, the crystal structure of the human lamin-B1 coil 2 segment (PDB ID: 3TYY[23]) was selected as the best template for homology modeling. The partial 3TYY structure was optimized and used as a template to generate 50 homology models using the Build Homology Models protocol. Probability Density Function total energy and Discrete Optimized Potential Energy scores were used to select the best model. Protein stability for mutants was calculated using the Calculate Mutation Energy (Stability) protocol. All molecular modeling and simulation were performed with Discovery Studio, v. 4.1, from BIOVIA (Accelrys Inc. San Diego, USA) with the default parameter setting.

### Genome editing by CRISPR/Cas9 in mouse embryos

To generate mice carrying the p.Arg587Trp disease-associated missense variant, we edited the Cchcr1 codon sequence of amino acid 591 in mice using the following protocol (Fig.3). Cas9 mRNA was prepared using a pBGK plasmid, as previously described.[67] The plasmid was linearized with XbaI and used as the template for *in vitro* transcription with an mMESSAGE mMACHINE T7 ULTRA kit (Ambion, Foster City, CA, USA). Single guide RNAs (sgRNAs) (guide1: GCTGTGTCAGCTCCTGACGG[AGG], guide2: GGAAGCTGCCAGCCTCCGTC[AGG]) were designed using CRISPR design (CRISPR.mit.edu). The templates for sgRNA synthesis were PCR amplified with primer sets (5’-TAATACGACT CACTATAGGGCTGTGTCAGCTCCTGACGGGTTTTAGAGCTAGAAATAGCA AG-3’ / 5’-AAAAAAAGCACCGACTCGG-3’ for sgRNA1, 5’-TAATACGACTCACTATAGGGGAAGCTGCCAGCCTCCGTCGTTTTAGAGCTAGAAATAGCAAG-3’ / 5’-AAAAAAAGCACCGACTCGG-3’ for sgRNA2) using pUC57-sgRNA vector (Addgene number: #51132) as a template. Then, 400 ng of gel-purified PCR products were subjected to RNA synthesis with a MEGAshortscript T7 Kit (Ambion), according to the manufacturer’s instructions.[68] Both Cas9 mRNA and sgRNA were purified with a MEGAclear kit (Ambion) and filtered by passing through an Ultrafree-MC filter (HV; 0.45 μm pore size; Millipore, Billerica, MA) to avoid clogging during microinjection. The single-stranded oligodeoxynucleotide (ssODN) (5’-CAGCAGTTGGAGGCAGCACGTCGGGGCCAGCAGGAGAGCACGGAGGAAGCTGCCAGC CTCtGgCAGGAGCTGACACAGCAGCAGGAAATCTACGGGCAAGGTGTGGGGGCGTGGCGGTGTGTG-3’) was synthesized by IDT (Coralville, IA, USA). C57BL/6N strain mouse zygotes were obtained using *in vitro* fertilization. One-cell stage fertilized mouse embryos were injected with 10 ng/μl of Cas9 mRNA, 10 ng/μl of sgRNA, and 20 ng/μl of ssODN. Injected eggs were cultured overnight in KSOM medium and the resulting two-cell embryos were transferred into the oviducts of pseudo-pregnant ICR females, as previously described[68].

### Mice

C57BL/6N mice were purchased from CLEA (Shizuoka, Japan) and maintained under specific pathogen-free conditions. Wild-type mice (8-to 10-month-old females and males) were used as control group and/or for calibration. This study used 27 mice (8- to 12-month-old females and males) carried the mutation generating amino-acid substitution. Randomization and blinding tests were not performed in this study. All animal procedures were done according to protocols approved by the Institutional Animal Care and Use Committee of Tokai University.

### Genotyping allele of target locus for generated mice with alkaline lysis method

Mouse tissue obtained by ear punch was added to 180 μl of a 50-mM NaOH solution and incubated at 95°C for 10 minutes. The lysate for PCR was obtained by neutralizing with 20 μl of 1 M Tris-HCl (pH 8.0) and centrifugation. PCR was performed in a reaction volume of 10 μl containing 1 μl of lysate, 0.2 U of KOD FX Neo (TOYOBO), 5 μl of 2× PCR buffer, 2 μl of dNTP (2 mM each), and 0.2 mM (final concentration) of each primer. Forward (AGCTGAGTGCCCACCTGAT) and reverse (TGTGTCTCAGTGCTGCCTTC) primers were used for sequencing (Greiner Bio-one). The thermal cycling profile was as follows: initial denaturation at 94°C for 2 minutes, then 35 rounds of amplification at 98°C for 10 seconds and 60°C for 25 seconds. The protocol for sequencing was performed as previously described in as above in ‘Variant discovery and genotyping of *CCHCR1* by Sanger sequencing’.

### RNA isolation

Total RNA was isolated from sections of mouse skin using ISOGEN (Nippon Gene, Tokyo, Japan), according to the manufacturer’s protocol, and treated twice with TURBO DNase (Ambion) to eliminate contaminating DNA. RNA was quantified using a NanoDrop 2000 (Thermo Fisher Scientific) and the quality of the extracted RNA was evaluated with a Bioanalyzer 2100 (Agilent Technologies).

### Microarray analysis

Fluorescent cRNA synthesis derived from skin RNAs was performed using a Low RNA Input Linear Amplification kit (Agilent Technology) and subjected to DNA microarray analysis with single-color microarray-based gene-expression analysis (SurePrint G3 Mouse GE, v. 2.0, 8×60 K, Agilent Technology). All procedures were performed according to the manufacturer’s instructions. Data from samples that passed the QC parameters were subjected to 75th percentile normalization and analyzed using Genespring GX (version 12, Agilent Technologies). Gene ontology enrichment analysis was performed with DAVID 6.8 (https://david.ncifcrf.gov/) using 3 functional database (Cellular Component, Biological Process and Molecular Function). Significance values were calculated between the AA and wild-type mice using Spearman’s rank correlation coefficient (paired, two-tailed).

### Quantitative PCR

cDNAs were synthesized using 1 μg of total RNA in a 20-μl total volume using SuperScript^®^ VILO^TM^ MasterMix (Thermo Fisher Scientific) and random hexamers. Quantitative PCR was performed using a StepOnePlus™ Real-Time PCR System, TaqMan^®^ Universal Master Mix, and TaqMan gene expression assay (Thermo Fisher Scientific), according to the manufacturer’s protocol. The primer-probe sets were as follows: Mm00652053_g1 (*Krt34*), Mm02345064_m1 (*Krt73*), Mm04208593_s1 (*Krtap3-3*), Mm04336701_s1 (*Krtap16-1*), Mm00478075_m1 (*Padi3*), Mm01214103_g1 (*S100a3*), Mm00461542_m1 (*Cchcr1*), and Mm99999915_g1 (*Gapdh*) (Thermo Fisher Scientific). All PCR reactions were performed in triplicate. Relative quantification of gene expression was performed using the 2^-ΔΔCt^ method[39]. Fold change values were calculated using *Gapdh* as the internal control and a dorsal skin sample from a wild-type mouse was used as a calibrator. Significance values were calculated between the AA and wild-type mice using Student’s t-test (unpaired, two-tailed).

### Immunohistochemistry

Anti-CCHCR1 (rabbit polyclonal) and anti-hair cortex cytokeratin antibodies (mouse monoclonal [AE13]) were obtained from Novus Biologicals (NBP2-29926) (Littleton, CO, USA) and Abcam (ab16113) (Cambridge, MA, USA), respectively. This CCHCR1 antibody is generated from rabbits immunized with a KLH conjugated synthetic peptide between 599-627 amino acids from the central region of human CCHCR1. The homology of the 29 peptides between human and mouse is 97% (28/29).

Deparaffinized skin sections were boiled in 10 mM of citrate buffer (pH 5.0) for antigen unmasking. Sections were incubated in Blocking One Histo (Nacarai tesque, Kyoto, Japan) for 30 minutes at room temperature, then incubated with primary antibodies [anti-CCHCR1 (1:100), anti-hair cortex Cytokeratin (1:200)] in PBS containing 5% Blocking One Histo and 0.05% Triton-X 100 overnight at 4°C. Sections were then washed and incubated with secondary antibodies [anti-mouse IgG-Alexa488 (1:500), anti-rabbit IgG-Alexa594 (1:500)] for 2 hours at room temperature. Controls for all immunostaining were simultaneously performed by omitting the primary antibody. Sections were cover-slipped using Vectashield with 4’,6-diamidino-2-phenylindole dihydrochloride (DAPI) (Vector Laboratories, Burlingame, CA, USA) for nuclei counterstaining and analyzed with Keyence BZ X-700 (Keyence, Tokyo, Japan).

### Morphology of hair shafts

Hairs were plucked from mice and analyzed by SEM using a JSM 6510LV (Jeol Co., Tokyo, Japan).

**Additional files**: Supplementary figures, tables and sequences.

**Figure S1**. Evaluation of pair-wise LD between 23 multi-allelic loci in MHC region. **Figure S2**. Evaluation of 16 variants extracted from all variants detected using NGS. **Figure S3** Sanger sequencing confirmation of rs142986308. **Figure S4** Schematic overview of CCHCR1 gene structure and variants with amino acid substitution. **Figure S5** Coiled-coil structure prediction of CCHCR1 in 26 haplotypes using COILS, v. 2.2. **Figure S6** Coiled-coil structure prediction of CCHCR1 in 5 selected haplotypes using Paircoil2. **Figure S7** Fixed hair loss area in representative AA mouse after onset of hair loss. **Figure S8** Hair loss area recovered in representative AA mouse. **Figure S9** Scatter plot between dorsal and ventral skin areas in same individual. **Figure S10** Scatter plots of comparisons between AA and wild-type skin. **Figure S11** Filtering scheme used for microarray analysis. **Figure S12** Analysis of expression in mouse skin using quantitative PCR and a comparative CT method. **Figure S13** Gene expression trends in skin biopsies from AA mouse 01 and 02. **Figure S14** Pair-wise LD between 24 variants with amino acid substitution in *CCHCR1*. **Figure S15** Analysis of pair-wise LD between 24 loci for investigating LD. **Figure S16** Quantification and verification of genomic libraries. **Table S1** Allelic association analysis of 23 loci spanning 2.45 Mb in the MHC region. **Table S2** Allelic association analysis of *HLA-C* gene. **Table S3** Overview of sequencing output by NGS. **Table S4** Filtering of variants identified by MHC region sequencing. **Table S5** Heterozygous variants identical in individuals with risk haplotype. **Table S6** Overview of 16 variants identical in 5 individuals with risk haplotype. **Table S7** SNV discovery and allelic association with *CCHCR1*. **Table S8** Homology of amino acid sequence to Hap26 in various species. **Table S9** List of 265 probes showing ≥2-fold up- or down-regulation in AA mice as compared to wild type. **Table S10** Inverse correlation of expression fold change values between Hoxc13- and AA mice. **Table S11** Enrichment analysis by DAVID in 246 genes showing ≥2-fold change in gene expression. **Table S12** Microsatellites and the primers. **Table S13** Primer sets for PCR direct sequencing in all exons of *CCHCR1* gene. Supplementary sequences Amino acid sequences were generated from haplotype nucleotide sequences (Table 2) as transcript variant3 (NM_019052) (Additional file1: Figure S4).

### List of abbreviations

Alopecia areata (AA), the major histocompatibility complex (MHC), the coiled-coil alpha-helical rod protein 1 (*CCHCR1*), keratin-associated proteins (KRTAPs), the human leukocyte antigen (HLA), linkage disequilibrium (LD), extended haplotype homozygosity (EHH), peptidyl arginine deaminase type III (*Padi3*), S100 calcium binding protein A3 (*S100A3*), trichohyalin (*Tchh*), homeobox C13 (*Hoxc13*), inner root sheath (IRS), outer root sheath (ORS), forkhead box N1 (*Foxn1*), the Database for Annotation, Visualization and Integrated Discovery (DAVID), and quantitative PCR (qPCR)

### Declarations

#### Ethics approval and consent to participate

Upon approval of the experimental procedures from the relevant ethical committees of Juntendo University (reference number:2013097) and Tokai University (reference number: 131-07), we obtained informed consent from all unrelated AA and healthy individuals prior to collection of DNA samples. All animal procedures were done according to protocols approved by the Institutional Animal Care and Use Committee of Tokai University.

### Consent for publication

Not applicable.

## Availability of data and materials

Complete DNA microarray data set is deposited at the Gene Expression Omnibus (GEO) database under GSE100630 (https://www.ncbi.nlm.nih.gov/geo/info/linking.html). To whom correspondence and material requests should be addressed: 143 Shimokasuya, Isehara, Kanagawa, 259-1193, Japan Tel: +81 463931121; Fax: +81 463964137;

## Authors’ contributions

A.Oka and S.I. conceived the project. A.Oka, M.O., and S.I. designed the research. A.T., E.K., and S.I. were involved in sample collection and clinical interpretation. A.Oka, K.H., S.S., N.M., T.H., M.K., H.M., M.O., Y.H., A.Otomo, S.H., and T.M. performed laboratory experiments. A.Oka, M.T.U., S.N., M.T., T.K., and S.M. contributed to the data analysis and statistical support. A.Oka, M.O., A. Otomo, S.H., H.I., S.B., and S.I. wrote the manuscript with contributions from all coauthors.

## Acknowledgments

We express our gratefulness to Wang Ting, Hisako Kawada Masayuki Tanaka, and Hideki Hayashi of the Support Center for Medical Research and Education, Tokai University.

